# Generation of salivary glands derived from pluripotent stem cells via conditional blastocyst complementation

**DOI:** 10.1101/2023.11.13.566845

**Authors:** Junichi Tanaka, Akihiro Miura, Yuko Shimamura, Youngmin Hwang, Dai Shimizu, Yuri Kondo, Anri Sawada, Hemanta Sarmah, Zurab Ninish, Kenji Mishima, Munemasa Mori

## Abstract

Various patients suffer from dry mouth due to salivary gland dysfunction. Whole salivary gland generation and transplantation is a potential therapy to resolve this issue. However, the lineage permissible to design the entire salivary gland generation has been enigmatic. Here, we discovered Foxa2 as a lineage critical for generating a salivary gland via conditional blastocyst complementation (CBC). Foxa2 linage, but not Shh nor Pitx2, initiated to label between the boundary region of the endodermal and the ectodermal oral mucosa before primordial salivary gland formation, resulting in marking the entire salivary gland. The salivary gland was agenesis by depleting Fgfr2 under the Foxa2 lineage in the mice. We rescued this phenotype by injecting donor pluripotent stem cells into the mouse blastocysts. Those mice survived until adulthood with normal salivary glands compatible in size compared with littermate controls. These results indicated that CBC-based salivary gland generation is promising for next-generation cell-based therapy.

## Introduction

Cell-based therapy for generating salivary glands is a promising next-generation therapy for patients suffering from dry mouth, rampant caries, and fungal infections due to Sjögren’s syndrome or the side effects of radiotherapy for head and neck cancers^1^. The strategies of isolating and expanding tissue-specific stem cells in the salivary gland tissues and their injection have been proposed ^2–5^, while the extent to which endogenous tissue stem cells with the ability to restore salivary gland function remain in damaged salivary glands remains controversial. To fill the gap of knowledge, we reported a strategy to transplant exogenous salivary gland progenitor organoids induced to differentiate from pluripotent stem cells (PSC)^6,7^. These PSC-derived salivary gland organoids were engrafted orthotopically into salivary gland-resected mice. However, post-transplantation, PSC-derived salivary gland organoids size was smaller than usual, and the recovery effect on salivary secretion was limited ^6^. A better strategy that produces whole salivary glands derived from PSCs to overcome size issues was needed.

Blastocyst complementation (BC) has been proposed as a promising option for creating fully functional organs derived from PSCs *in vivo*^8^. This unique technique has further evolved into generating intra- and interspecies organs such as kidneys, pancreas, and blood vessels ^9–12^. Previously, we refined the BC approach to the conditional blastocyst complementation (CBC) strategy, which allowed us to generate fully functional organs by targeting a specific lineage to be rescued, followed by PSC injection^13^. Although developing an emptying niche in a lineage is the key to a successful CBC, the lineage and the genes that vacate the salivary gland niche have been unknown.

It is well-known that a Fgf10 ligand, expressed in mesenchymal cells during mouse and human development, is crucial for salivary gland organogenesis through interacting with its receptor, Fgfr2, expressed in the epithelial cells^14,15^. In the systemic knockout (KO) mice of Fgfr2, various organs, including salivary glands, lungs, and limbs, show aplasia or hypoplasia phenotypes^16–19^, while the timing and lineage that requires Fgfr2 for salivary gland formation have been unclear.

The origin of the salivary glands before primordial salivary gland initiation has been debated. Based on previous *Sox17-LacZ* lineage tracing mouse analysis studies, most endodermal Sox17-lineage does not label salivary gland epithelium^20^. Thus, the salivary gland has been believed to originate from the ectodermal oral mucosa but not the endoderm. Contradictorily, Sonic Hedgehog (Shh), primarily known as a marker of the endodermal epithelium, has also been reported to label salivary gland epithelial lineage^21^. This confusion arises because the developmental origin of the submandibular and sublingual glands anatomically arises around the boundary region between the ectodermal and the endodermal-derived oral epithelium (OE)^22,23^. Based on this, for generating PSCs-derived salivary glands by CBC, we investigated the lineage to label the boundary region, which is potentially critical for salivary gland formation.

## Results

### Foxa2 lineage contributes to salivary gland development

Shh lineage has been reported to contribute to endoderm, pharyngeal pouch^24^, and salivary gland epithelium at the later stage of development at E15.5, while Shh regulates the branching morphogenesis of salivary glands^25^. On the other hand, the Pitx2 lineage is another potential lineage of the salivary gland because its protein is known to be expressed in embryonic oral and dental epithelium before the salivary gland formation, and the Pitx2 defect results in a range of developmental deficits, including defective body-wall closure, right pulmonary isomerism, and altered cardiac position^26–28^. Using *Shh^Cre/+^; Rosa^LSLtdTomato/+^* or *Pitx2^Cre/+^; Rosa^LSLtdTomato/+^* lineage-tracing mice, we examined whether Shh or Pitx2 lineage could contribute to salivary gland epithelium. We found that 97.2% ± 1.59 in Shh lineage tracing mice and 98.7% ± 0.66 in Pitx2 lineage tracing mice of the salivary gland epithelium was tdTomato positive E18.5 in both of the lineage tracing analyses (Figure S1A, S1B, and S1C), supported by the previous studies^21^. Surprisingly, we found that the salivary gland was usually formed in both lineage-specific Fgfr2 KO mice (Figure S1A and S1B). These results indicate that the Pitx2 or Shh lineage-based strategy is insufficient for inducing the phenotype of salivary gland agenesis, while it has been unclear why Fgfr2 deficiency did not cause salivary gland defective phenotype in vivo despite the evidence that Fgfr2 is known to be critical for the salivary gland branching morphogenesis in ex-vivo culture and knockout study^14,29^.

To address this issue, we further investigated another potential lineage involved in the salivary gland precursor niche around the boundary region of endoderm and ectoderm by analyzing deposited single-cell RNA-seq (scRNAseq) data from E8.5 mouse^30^ and E12.0 OE^31^ because the organ agenesis phenotype caused by Fgfr2 deficiency in other organs, like lungs, requires to target the precursor niche but not the progenitors to obtain after the organ initiation^13,32^. In the E12.0 OE scRNAseq analysis, we identified high-expression genes in the posterior-lateral epithelium (PL) and tongue epithelium (T) adjacent to the salivary gland primordium (Figure S1D, S1E, and S1F). The endodermal marker Foxa2 showed high expression in T, while Sox9, Pitx2, and Shh expressed in salivary gland, anterior region, and tooth initial knot, respectively (Figure S1D and S1E)^20^.

Using E8.5 mouse scRNAseq, we identified Foxa2 as the potential gene expressed around the boundary region of E8.5 OE between ectoderm and endoderm, indicating the salivary gland precursors of the growing boundary region (black dotted area) for future salivary gland primordia formation (Figure S1G and S1H). Conversely, neither Shh nor Pitx2 was expressed on the precursor of the E8.5 OE (Figure S1H). These data suggested that the Foxa2 may label the boundary of the E10.5 OE as a salivary gland precursor niche before salivary gland primordia formation.

To confirm this observation, we performed lineage tracing studies utilizing *Foxa2^Cre/+^; Rosa^tdTomato/+^* mice. Before the boundary region formation by the connection of ectodermal OE and the endodermal epithelium (Figure 1A), we found that the Foxa2 lineage cells appeared a few in the ectodermal OE (Figure 1B). Significantly, the boundary region of the E9.5 epithelial cells on the oral floor was entirely labeled by the tdTomato-positive Foxa2 lineage (Figure 1C, arrowhead). In contrast, Shh or Pitx2 lineage did not mark tdTomato on the E9.5 OE of the boundary region (Figure S1I, arrows). At E10.5, the Foxa2 lineage labeling had extended to the anterior region of the OE, while it did not contribute to the ventral surface epidermis (VSE) (Figure 1D). Interestingly, at this point, the expression of the Foxa2 protein itself was limited to the posterior OE (Figure 1E). As evidenced by the tdTomato signal and histological analysis (Figure 1F and 1G), the Foxa2 lineage in the oral epithelium contributed to the invaginating OE (Figure 1G, arrow) and primordium formation (Figure 1G, asterisk) at E13.5 and occupied the adjacent epithelium. Remarkably, we observed that the Foxa2 lineage significantly labeled 99.3% ± 0.64 of salivary E15.5 gland epithelium compared to mesenchyme labeling (2.9% ± 2.11) (Figure 1H and 1I).

**Figure 1.**
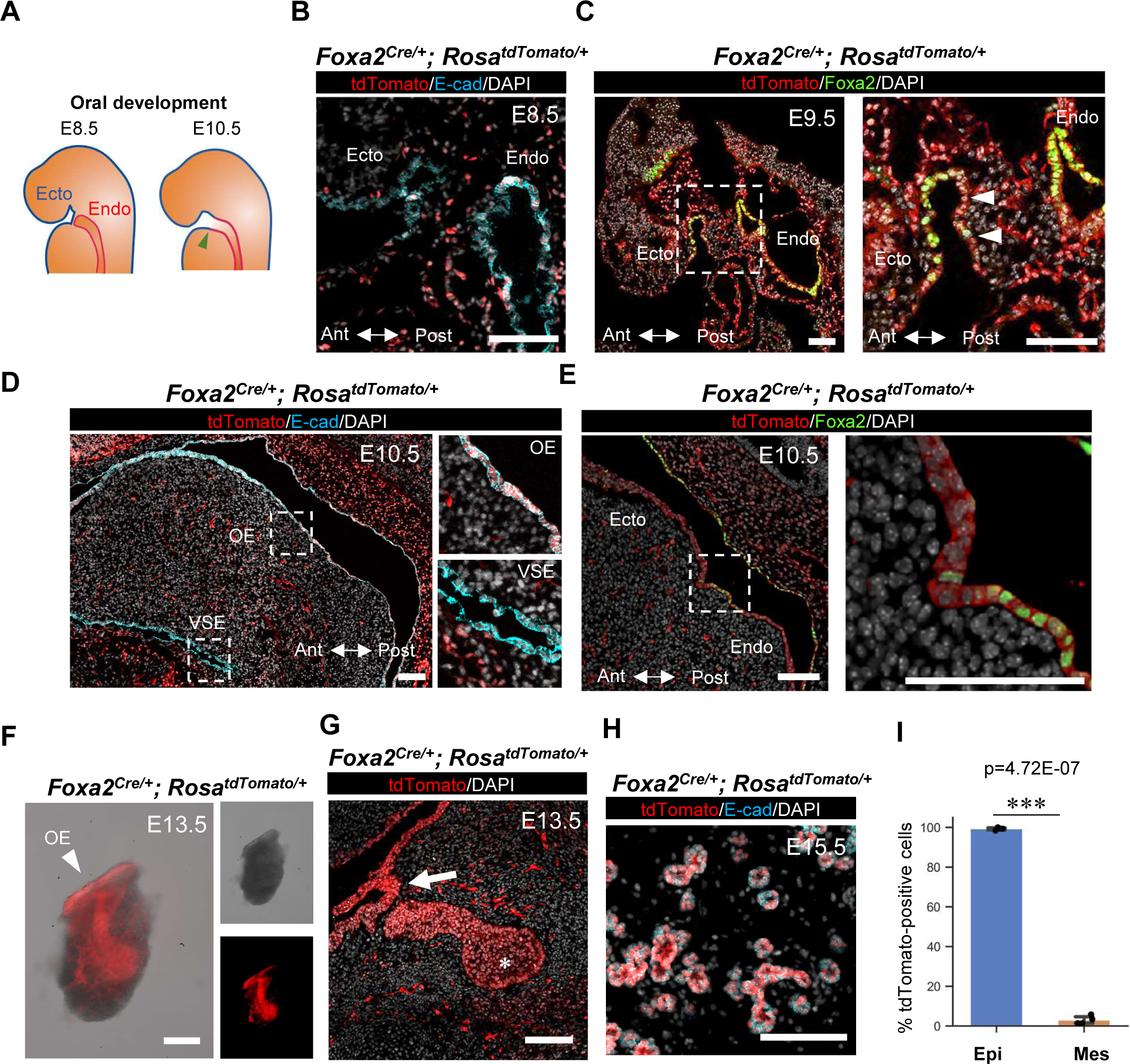
Foxa2 lineage labeled boundary region between endoderm and ectoderm, leading to mark whole salivary gland epithelium. **(A)** Schematics of the boundary region development located between endodermal (a red line) and ectodermal oral (a blue line) epithelial cells at embryonic day (E) 8.5 and E10.5 before salivary gland primordial formation. The green arrowhead indicates the boundary region of salivary gland development. Ecto: Ectoderm, Endo: Endoderm. **(B-E)** Representative immunofluorescence (IF)-confocal imaging of the ectodermal (Ecto) oral mucosal and endodermal (Endo) epithelial boundary in E8.5 **(B)**, E9.5 **(C)**, and E10.5 **(D, E)** *Foxa2^Cre/+^; Rosa^LSL-tdTomato/+^* lineage-tracing mice. Immunostaining of tdTomato: red, E-cadherin (E-cad): cyan, DAPI: gray, and Foxa2: green. Each right panel of **C-E**: an enlarged image of a white dotted box. Ant: arteriolar, Post: posterior axis, VSE: ventral surface epidermis, OE: oral epithelium. (N=3 at each time point, biological replicates) **(F)** Representative merged image (left) of bright-field (right top) and tdTomato fluorescent signal (right bottom) of the isolated salivary gland from E13.5 *Foxa2^Cre/+^; Rosa^LSL-tdTomato/+^* lineage tracing mice. (N=3) **(G, H)** Representative confocal imaging of E13.5 salivary gland primordium **(G)** and E15.5 salivary glands **(H)** *Foxa2^Cre/+^; Rosa^LSL-tdTomato/+^* lineage-tracing mouse. (N=4) **(I)** Graphs: the morphometric lineage-tracing analysis: % of Foxa2-lineage labeling in E-cad-positive epithelial (blue bar) and E-cad-negative mesenchymal (orange bar) cells from E15.5 *Foxa2^Cre/+^; Rosa^LSL-tdTomato/+^* lineage tracing mouse salivary glands (N=4). Statistical analyses: unpaired Student’s t-test, significant: p<0.05. ***p < 0.001. Error bars represent mean ± SD. Scale bars: 100 μm.

### Foxa2-driven Fgfr2 conditional knockout (cKO) caused salivary gland agenesis phenotype

Since the Foxa2 lineage contributes to the E8.5∼E9.5 boundary region between endodermal and ectodermal OE, leading to the entire salivary gland labeling, we investigated whether mice with Fgfr2 depletion in the Foxa2 lineage would show salivary gland agenesis phenotype. We utilized the *Foxa2^Cre/+^; Fgfr2^flox/flox^; Rosa^LSL-tdTomato/+^* mice. This mouse model, which we previously reported, exhibits an absence of lung formation^32^. In the Fgfr2 heterozygous knockout mice (*Foxa2^Cre/+^; Fgfr2^flox/+^; Rosa^LSL-tdTomato/+^*, hereafter, *Fgfr2^hetero^*), tdTomato-positive salivary gland primordia were detected at the base of the oral cavity at E13.5 (Figure 2A). Conversely, tdTomato-positive tissue was absent in *Fgfr2^cKO^*mice (*Foxa2^Cre/+^; Fgfr2^flox/flox^; Rosa^LSL-tdTomato/+^*), suggesting the salivary gland agenesis phenotype (Figure 2B). In contrast to the lineage tracing results (Figure 1G), histological analysis showed that tdTomato^+^ OE invagination into mesenchyme (Figure 2B, arrow), but it lost salivary gland primordia formation in the *Fgfr2^cKO^* mice at 13.5 (Figure 2B, asterisk). This result indicates that Fgfr2 is critical for initiating salivary gland primordia formation after the invagination of oral epithelium but not essential for invagination.

**Figure 2.**
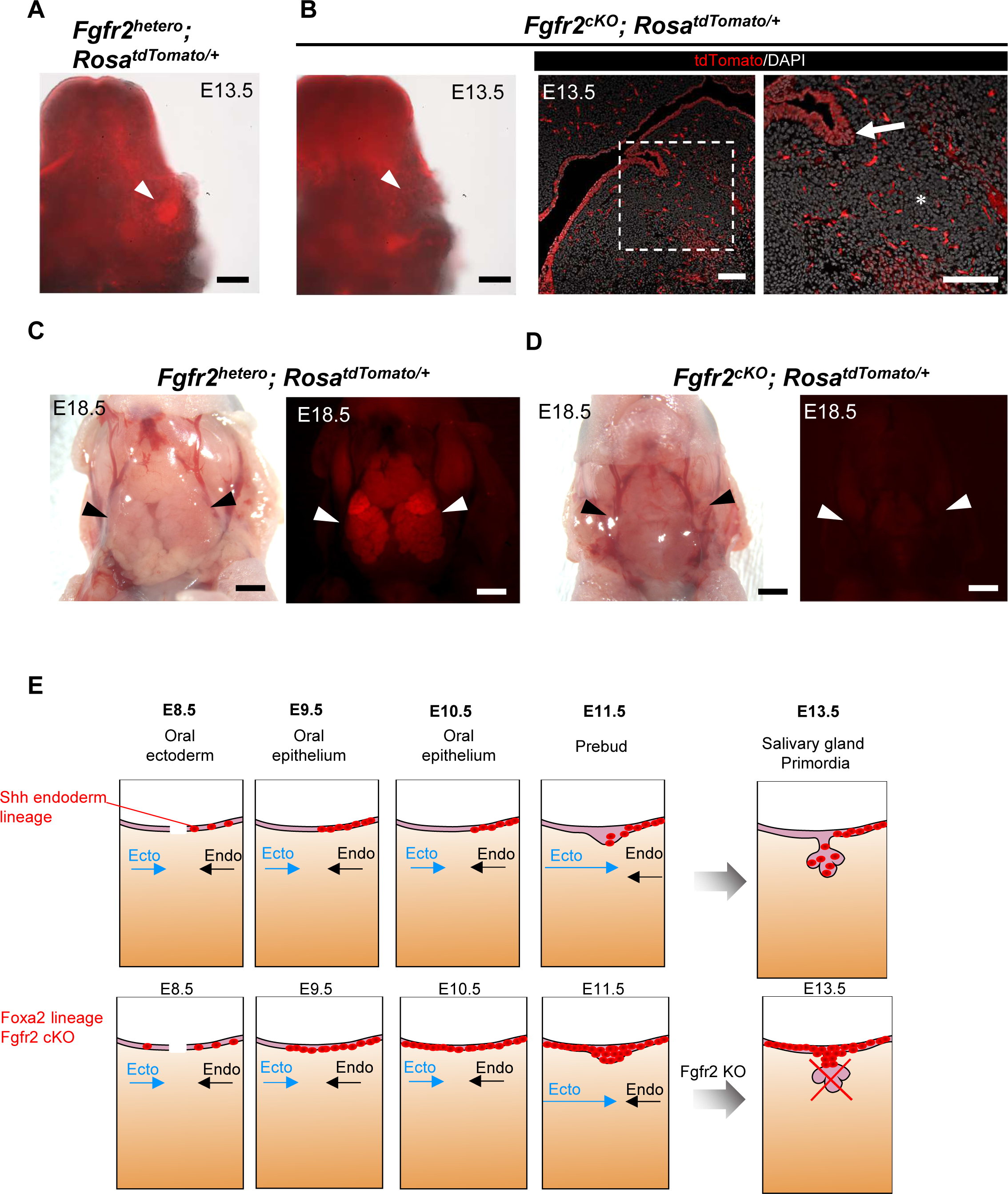
Foxa2-driven Fgfr2 conditional knockout (*Fgfr2^cKO^*) caused salivary gland agenesis phenotype. **(A, B)** Representative macroscopic merged images of bright-field and tdTomato fluorescent signals of salivary gland primordiu formation (white arrowheads) from E13.5 *Fgfr2^hetero^* (*Foxa2^Cre/+^; _Fgfr2_flox/+_; Rosa_LSL-tdTomato/+*_) mice **(A)** and E13.5 *Fgfr2*_*cKO* _(*Foxa2*_*Cre/+_; Fgfr2_flox/flox_; Rosa_LSL-tdTomato/+*_):_ No salivary gland primordia formation (**B**, left). Representative IF-confocal imaging of the salivary gland rudiment in E13.5 *Fgfr2^cKO^* (*Foxa2^Cre/+^; Fgfr2^flox/flox^; Rosa^LSL-tdTomato/+^*) mice (N=3): No salivary gland primordium formation (asterisk) but invaginating OE occurs (arrow), indicating the salivary gland rudiment **(B,** middle and right). **(C, D)** Representative macroscopic bright-field images (left) and tdTomato fluorescent signals (right) of salivary glands (arrowhead) from E18.5 *Fgfr2^hetero^* (*Foxa2^Cre/+^; Fgfr2^flox/+^; Rosa^LSL-tdTomato/+^*) **(C)** and E18.5 *Fgfr2^cKO^* (*Foxa2^Cre/+^; Fgfr2^flox/flox^; Rosa^LSL-tdTomato/+^*) **(D)** (N=3): Salivary gland agenesis phenotype in the E18.5 *Fgfr2^cKO^* (*Foxa2^Cre/+^; Fgfr2^flox/flox^; Rosa^LSL-tdTomato/+^*). tdTomato: red. **(E)** Proposed schematic models for the initial salivary gland development from the junction of ectodermal oral mucosa and endodermal epithelium based on Foxa2 or Shh lineage tracing analysis: When ectodermal oral mucosa (Ecto: blue arrows) and endodermal (Endo: arrows) epithelium is closer and connecting, Shh lineage (red) labeled cells appeared only on the oral endodermal side (Endo: black arrows) of the E8.5 and E9.5 boundary region but not oral ectodermal side of salivary gland precursors (SGP) before the boundary formation. Later, the Shh lineage labeled nearly the entire E15.5 epithelium, supported by a previous study^21^. In contrast, the Foxa2 lineage initiated labeling the E8.5 SGP and the entire E9.5 SGP on the boundary region before the invagination of the salivary gland precursors. Therefore, the knockout of Fgfr2 in the Foxa2 lineage resulted in the invagination of SGP but failed to form salivary gland primordia, leading to the salivary gland agenesis phenotype. Scale bars: B, left: 200 μm, B, right: 100 μm, C, D: 1mm.

To ensure that these observations were not due to developmental delays by the Fgfr2 deficiency in the Foxa2 lineage, we performed the macroscopic analysis at E18.5 *Fgfr2^cKO^* mice. While tdTomato-positive salivary gland tissues were formed in the E18.5 *Fgfr2^hetero^*mice (Figure 2C, arrow), tdTomato-positive salivary glands were not observed in the *Fgfr2^cKO^*mice (Figure 2D, arrow). These results indicate the salivary gland agenesis phenotype rather than a developmental delay. Taken together, Fgfr2 depletion in the Foxa2 lineage-based niche is efficient for vacating the niche critical for salivary gland formation, which can be utilized for CBC (Figure 2E).

### Conditional Blastocyst Complementation using Foxa2-driven Fgfr2 cKO mice rescued the salivary gland formation during development

To address CBC would rescue the salivary gland agenesis phenotype, we injected donor mouse iPSCs expressing nuclear EGFP (nGFP) into host *Fgfr2^cKO^* mice. In this CBC approach, donor iPS cell-derived cells were GFP-positive, while host cells from the Foxa2 lineage were tdTomato-positive, and non-Foxa2 lineage cells were tdTomato-negative, allowing to distinguish visually (Figure 3A). Analysis at E17.5 revealed the formation of chimeric mice expressing both GFP and tdTomato signals. Furthermore, no significant external abnormalities were observed in the *Fgfr2^cKO^*, nEGFP^+^ iPSC chimeric mice, as well as littermate *Fgfr2^hetero^*, nEGFP^+^ iPSC chimeric mice (Figure 3B and 3C). Macroscopic analysis showed sporadic GFP and tdTomato signals were detected in the *Fgfr2^hetero^* chimeric mice’s submandibular and sublingual glands (Figure 3B). In contrast, in the salivary glands of *Fgfr2^cKO^*, nEGFP^+^ iPSC chimeric mice, a strong GFP signal was observed, while the tdTomato signal was low (Figure 3C). To quantify this observation, we performed a histological analysis. In the *Fgfr2^hetero^*chimeric mice, the E-cadherin (E-cad) positive salivary gland epithelium was formed as a chimera of host tdTomato-positive cells and donor nGFP-positive cells (Figure 3D). In the *Fgfr2^cKO^*, nEGFP^+^ iPSC chimeric mice, all the E-cad-positive salivary gland epithelium was rescued exclusively by donor nEGFP-positive cells without tdTomato^+^ host cells (Figure 3E). This chimerism rate in the *Fgfr2^cKO^*, nEGFP^+^ iPSC chimeric mice was significantly higher than in the heterozygous, nEGFP^+^ iPSC chimeric mice (*Fgfr2^cKO^*, nEGFP^+^ iPSC vs. *Fgfr2^htetero^*, nEGFP^+^ iPSC: 100.0% ± 0 vs. 74.7% ± 13.44) (Figure 3F). In contrast, the chimerism in mesenchymal cells did not show a significant difference between the *Fgfr2^hetero^* and *Fgfr2^cKO^* chimeric mice (*Fgfr2^cKO^*, nEGFP^+^ iPSC vs. *Fgfr2^htetero^*, nEGFP^+^ iPSC: 62.8% ± 16.35 vs. 64.3% ± 19.25) (Figure 3G).

**Figure 3.**
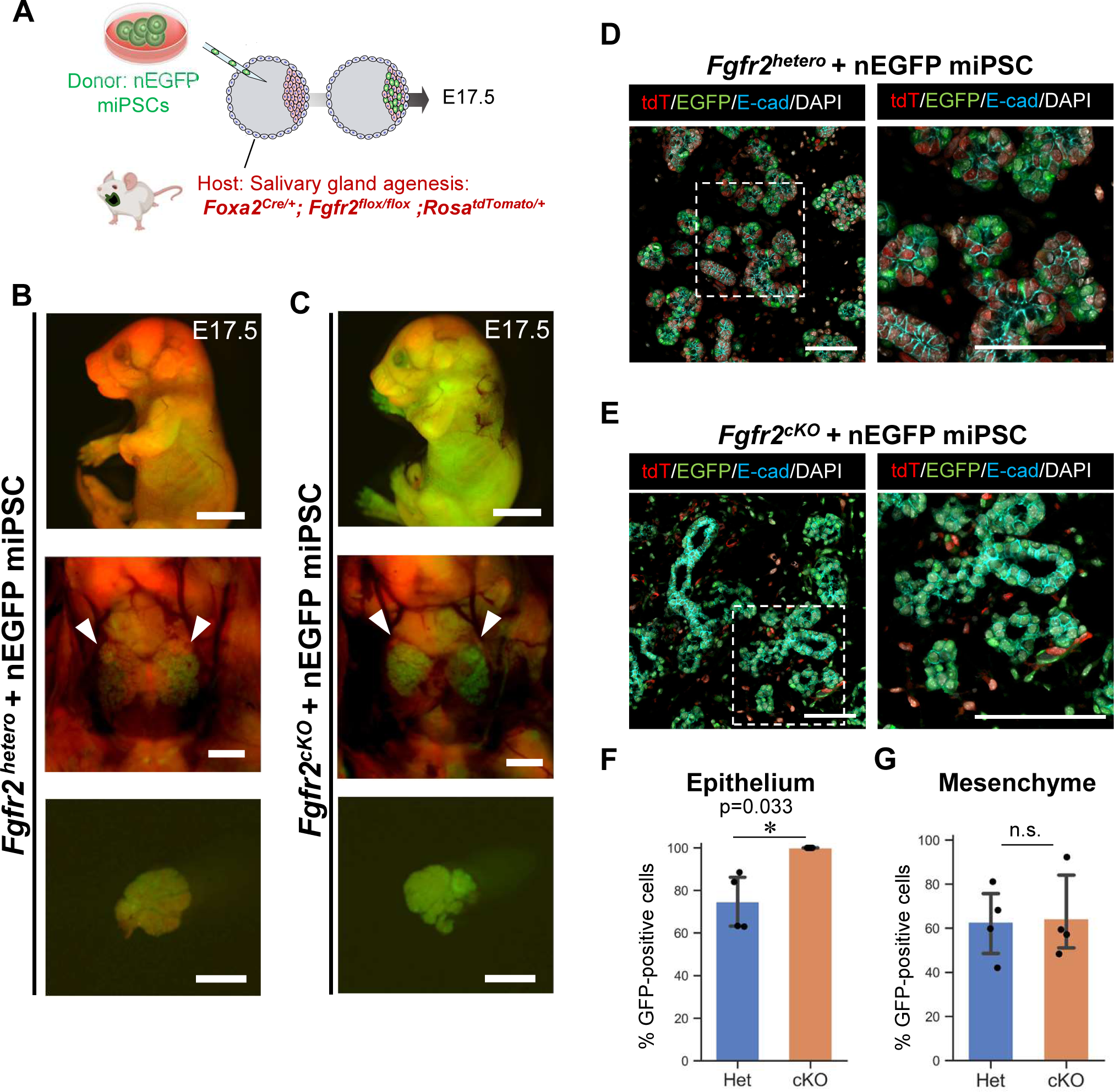
Rescue of salivary gland agenesis phenotype of *Fgfr2^cKO^* mice by Foxa2-lineage-based conditional blastocyst complementation (CBC) **(A)** Schema of CBC experiment: we injected nuclear EGFP (nEGFP)-positive mouse iPSCs (nEGFP miPSC) into the blastocysts of Foxa2-driven *Fgfr2^cKO^* (*Foxa2^Cre/+^; Fgfr2^flox/flox^; Rosa^LSL-tdTomato/+^*) mice or littermate control: *Fgfr2^hetero^* mice (*Foxa2^Cre/+^; Fgfr2^flox/+^; Rosa^LSL-tdTomato/+^*). **(B, C)** Representative images of fluorescent signals of the host (tdTomato: red) and donor nEGFP signal of nEGFP miPSC (green) from embryo (top), salivary glands (middle: arrows), and isolated salivary gland (bottom) from the E17.5 chimeric mice of *Fgfr2^cKO^*+nEGFP miPSC **(C)** or littermate control: *Fgfr2^hetero^*+nEGFP miPSC **(B)**. **(D, E)** Representative IF-confocal imaging of the salivary glands in the E17.5 chimeric embryos of *Fgfr2^cKO^*+nEGFP miPSC **(E)** or littermate control: *Fgfr2^hetero^*+nEGFP miPSC **(D)**. Immunostaining of tdTomato: red, EGFP: green, E-cadherin (E-cad): cyan and DAPI: gray. Each right panel of **D**, **E**: an enlarged image of a white dotted box. **(F)** graphs: the morphometric analysis: % of nGFP-positive donor cells in E-cad-positive epithelial cells from E17.5 chimeric embryos of *Fgfr2^hetero^*+nEGFP miPSC. (n = 3 per biological replicates, 5 fields per group). **(F, G)** The morphometric analysis: % of nGFP-positive donor cells in E-cadherin^+^ epithelium **(F)** or E-cadherin^-^ mesenchyme **(G)** from E17.5 chimeric embryos of *Fgfr2^cKO^*+nEGFP miPSC or littermate control: *Fgfr2^hetero^*+nEGFP miPSC. (N = 4 per biological replicates, 5 fields per group). Statistical analyses: unpaired Student’s t-test, significant: *p<0.05. no significant change: n.s. Error bars represent mean ± SD. Scale bars: B, C: 5 mm (top) and 1 mm (middle and bottom), D, E: 100 μm.

To confirm whether the fully complemented salivary glands in the *Fgfr2^cKO^*, nEGFP^+^ iPSC chimeric mice differentiate well, we examined the expression of various progenitor and differentiation markers in the complemented salivary glands (Figure S2). Indeed, E17.5 nEGFP^+^ intercalated duct cells and acinar cells expressed Sox9, a marker for salivary gland progenitors. nEGFP^+^ rescued ductal cells expressed Keratin 5 (K5), a marker for basal cells, and myoepithelial marker α-Sma around the acinar cells. E17.5 acinar cells expressed Mist1, a mature acinar marker, and Aqp5, a marker of mature acinar aggregated on the luminal surface of the Mist1^+^ acinar cells. These results indicate that the salivary gland epithelial cells complemented with mouse iPSCs develop and differentiate well at E17.5 without any developmental delays.

### Complemented salivary glands develop well enough in size and differentiation until adulthood

Based on this, we further examined whether the CBC-mediated rescued salivary glands would develop well until adulthood. To address this question, we injected donor mouse PSC^13^ expressing EGFP under CAG promoter (PSC^CAG-EGFP^) into the host Foxa2-driven Fgfr2 cKO mice. Based on the macroscopic fluorescent analysis of 4-week-old mice, the salivary glands of Fgfr2 heterozygous, PSC^CAG-EGFP^ chimeric mice formed a chimera consisting of tdTomato^+^ host cells and EGFP^+^ donor cells (Figure 4A). In contrast, the *Fgfr2^cKO^*, PSC^CAG-EGFP^ chimeric mice exhibited little tdTomato-positive signals (Figure 4B). In histological analysis, the salivary glands of the *Fgfr2^hetero^*, PSC^CAG-EGFP^ chimeric mice showed both tdTomato and EGFP signals in the epithelial cells (Figure 4C). Conversely, in the rescued *Fgfr2^cKO^*, PSC^CAG-EGFP^ chimeric mice, the epithelial compartment was entirely labeled by EGFP, while the mesenchymal cells showed a few tdTomato signals (Figure 4D). Furthermore, there was no significant difference in the weight of the complemented submandibular and sublingual glands between the chimeric heterozygous mice and the *Fgfr2^cKO^* mice (Figure 4E). This indicates that a sufficiently sized and functionally adequate salivary gland tissue was generated from donor PSCs via the Foxa2-lineage-based CBC approach. Additionally, we compared the EGFP^+^ donor cell chimerism with differentiation markers (the proportion of EGFP^+^ cells with each marker) between the *Fgfr2^cKO^*, PSC^CAG-EGFP,^ and *Fgfr2^htetero^*, PSC^CAG-EGFP^ chimeric mice (Figure 4F-H). In the chimeric salivary glands, GFP was labeled with the marker of Aqp5^+^ acinar cells (*Fgfr2^cKO^*, PSC^CAG-EGFP^ vs. *Fgfr2^htetero^*, PSC^CAG-EGFP^: 100.0% ± 0 vs. 82.1% ± 7.05), which aggregates on the luminal side with abundant cytoplasm, and of α-Sma^+^ spindle-shaped myoepithelial cells (*Fgfr2^cKO^*, PSC^CAG-EGFP^ vs. *Fgfr2^htetero^*, PSC^CAG-EGFP^: 100.0% ± 0 vs. 59.9% ± 9.75) surrounding the acini reflect the tissue structure of adult salivary glands as well as K5^+^ basal cells (*Fgfr2^cKO^*, PSC^CAG-EGFP^ vs. *Fgfr2^htetero^*, PSC^CAG-EGFP^: 100.0% ± 0 vs. 63.3% ± 13.39) (Figure 4H). Overall, in the complemented salivary glands in the *Fgfr2^cKO^*, PSC^CAG-EGFP^ mice, the Aqp5^+^ acinar cells, K5^+^ basal cells, and α-Sma^+^ myoepithelial cells all showed a GFP-positive rate of 100%, significantly higher than that in *Fgfr2^htetero^*, PSC^CAG-EGFP^ mice (Figure 4H). These results demonstrated that the emptying niche by depleting Fgfr2 in the Foxa2 lineage is critical for generating a fully-sized, well-differentiated adult salivary gland, followed by the PSC injection into the blastocysts.

**Figure 4.**
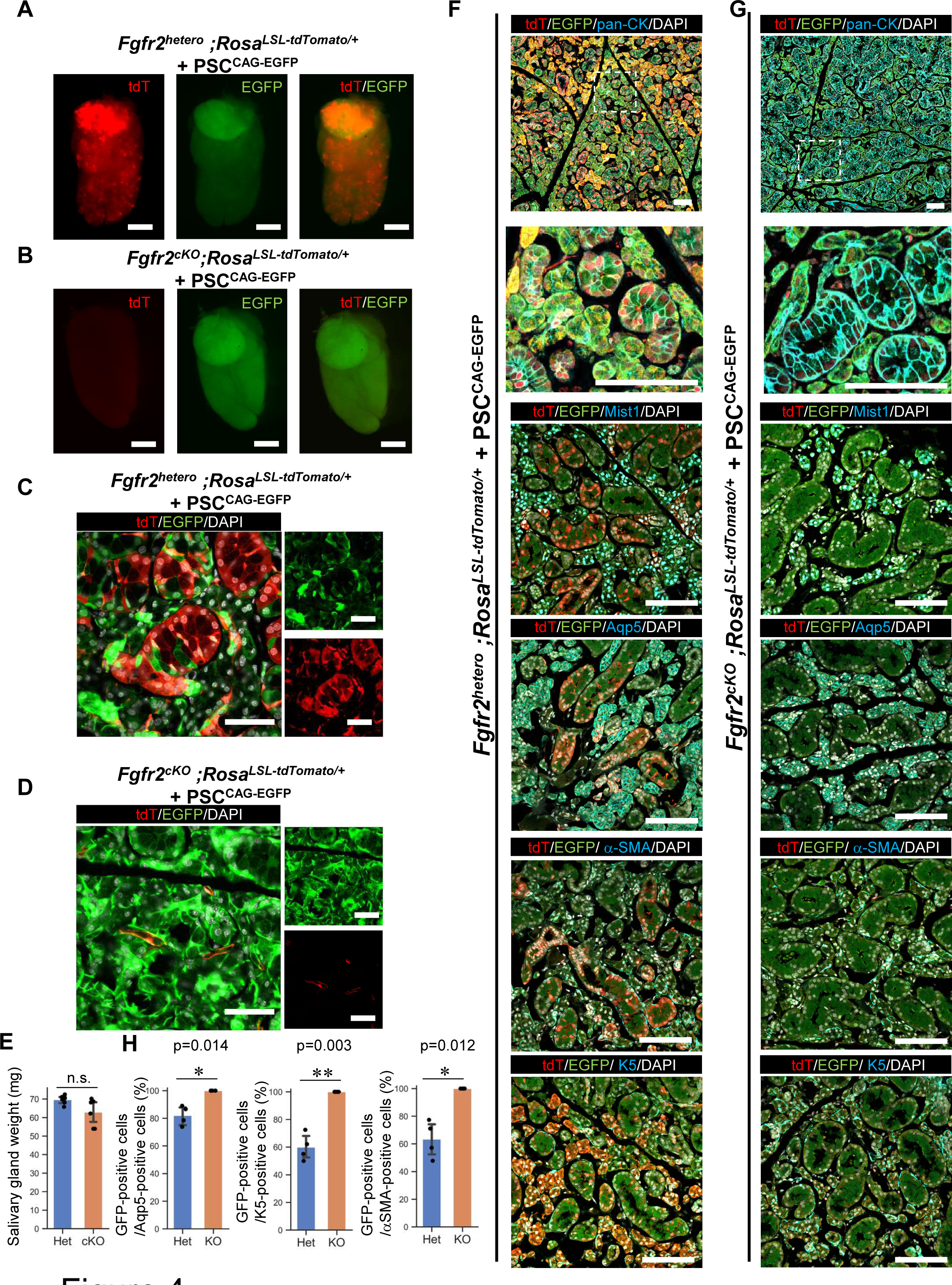
Fully mature adult salivary gland generation via Foxa2-lineage-based CBC. **(A, B)** Representative images of fluorescent signals of the host (tdTomato: red) and donor EGFP signal of PSC^CAG-EGFP^ (green) from isolated salivary gland from the 4 weeks chimeric mice of littermate control: *Fgfr2^hetero^*, PSC^CAG-EGFP^ **(A)** or *Fgfr2^cKO^*, PSC^CAG-EGFP^ **(B) (C, D)** Representative confocal imaging for native fluorescence signals of the salivary glands in the 4 weeks chimeric embryos of littermate control: *Fgfr2^hetero^*, PSC^CAG-EGFP^ **(C)** or *Fgfr2^cKO^*, PSC^CAG-EGFP^ **(D)**. **(E)** Analysis of salivary gland weight. The combined weight of the submandibular gland and the sublingual gland was measured. Het: *Fgfr2^hetero^*, PSC^CAG-EGFP^, cKO: *Fgfr2^cK^*, PSC^CAG-EGFP^. Statistical analyses: unpaired Student’s t-test, significant: p<0.05. no significant change: n.s. Error bars represent mean ± SD. **(F, G)** Representative IF-confocal imaging of the salivary gland from the 4 weeks chimeric mice of littermate control: *Fgfr2^hetero^*, PSC^CAG-EGFP^ **(F)** or *Fgfr2^cKO^*, PSC^CAG-EGFP^ **(G)**. The second panel from the top is an enlarged image of a white dotted box. Scale bars: 100 μm. **(H)** The morphometric analysis: % of EGFP-positive donor cells in Aqp5^+^ acinar cells (left), % of EGFP-positive donor cells in K5^+^ basal cells (middle), % of EGFP-positive donor cells in α-Sma^+^ myoepithelial cells (right) from the 4 weeks chimeric mice of littermate control: *Fgfr2^hetero^*, PSC^CAG-EGFP^ (Het) or *Fgfr2^cKO^*, PSC^CAG-EGFP^ (KO). (N=4 per biological replicates, 5 fields per group). Statistical analyses: unpaired Student’s t-test, significant: p<0.05. no significant change: n.s. *p<0.05. **p < 0.01. Error bars represent mean ± SD. Scale bars: A, B: 2 mm, D: 50 μm.

### Discussions

After radiotherapy for head and neck cancers^33–35^ or due to Sjögren‘s syndrome^36,37^, salivary gland acinar cells disappear by fibrosis, resulting in hyposalivation. Hyposalivation can cause various symptoms, including dry mouth, rampant caries, and fungal infections, but current treatment options for these symptomatic diseases are limited^38^. In this regard, cell-based therapy is one of the most promising approaches. We harnessed the blastocyst complementation (BC) approach to generate a fully functional salivary gland. The salivary glands produced by CBC showed weight equivalent to littermate control and well differentiation, indicating their promising potential as a source for future transplantation^39^.

Fgf7 and Fgf10, which are ligands of Fgfr2, have been proven to be crucial for the branching formation and duct elongation of salivary glands using *ex vivo* organ culture^14,40^. In this study, we provided new evidence that *Fgfr2^cKO^* in the Foxa2 lineage results in salivary gland agenesis, yet invaginating OE; salivary gland rudiment formation still occurs. Our results indicate that Fgfr2 is critical for the salivary gland primordia formation. We rescued this phenotype in the CBC experiment, and the salivary gland epithelium was entirely replaced by donor cells derived from PSCs. These results suggest that generating the vacant niche by Fgfr2 depletion in the invaginating OE was sufficient to rescue this phenotype via CBC, leading to complementing the entire salivary gland epithelium.

Salivary gland primordia arise from OE at E11.5. The ectodermal OE arises from the boundary region after the junction of the ectoderm and the endoderm around E8.5∼E9.5, but its exact origin has been unknown. Intriguingly, Shh lineage-tracing mice are known to label endoderm and also all salivary gland epithelial cells using *Shh^Cre/+^; Rosa^LSL-YFP/+^*mice, while the *Sox17 ^Cre/+^; Roa^LSL-LacZ/+^* mice, which is another endodermal lineage-tracing mice, do not label salivary gland epithelial cells^20,21^.

We revealed that Foxa2 is the lineage marking around the E9.5 boundary region critical for the salivary gland primordial formation. In the previous study using Krt14 lineage-driven Fgfr2 cKO, a small salivary gland primordium is formed^41^, while our lineage-tracing analysis using Shh or Pitx2 did not label the E9.5 boundary region of the oral mucosa. Based on these data, single-cell RNAseq data allowed us to draw the area most likely indicating the boundary region (black dotted lines in Figure S1H). These scRNAseq results indicated that targeting the boundary region using the Foxa2 lineage is critical for controlling the entire salivary gland epithelial precursor behaviors before primordial salivary gland formation, while extensive analyses of the boundary region at a single-cell level are required in the near future.

Systemic Fgfr2 knockout^40^ and our Foxa2-driven Fgfr2 knockout phenotype support the requirement of Fgfr2 for the salivary gland primordia formation. Our lineage tracing analysis using *Shh^Cre/+^; Rosa^tdTomato/+^* mice faithfully labeled endoderm but not OE right before the E8.5 boundary regions, while the previous study showed the lineage labels at E15.5 entire salivary gland^21^, and we showed E18.5 entire salivary gland, suggesting that the Shh lineage spontaneously increased its labeling from the boundary region formation to E15.5 entire salivary gland.

Since the Shh-driven Fgfr2 knockout phenotype showed lung agenesis phenotype, the Cre driver activity and genotyping cannot be mistaken in our experiments^13,32^. Interestingly, the Shh-driven Fgfr2 knockout phenotype did not show gross morphological changes in E18.5 salivary glands compared to the littermate controls, which is contradictory evidence of the Fgfr2’s effect in ex vivo culture^14^. Based on our data, we speculate that Fgfr2 depletion requires specific time windows to induce salivary gland agenesis phenotype, most likely in the salivary gland precursors around the boundary regions, as the lung agenesis model requires the depletion of the genes before the lung primordia formation^13,42^. For the exact requirement of Fgfr2 during salivary gland branching morphogenesis in vivo, spatiotemporal Fgfr2 depletion using *Shh^CreERT^*^2^*^/+^; Fgfr2^flox/flox^* or *Foxa2 ^CreERT^*^2^*^/+^; Fgfr2^flox/flox^* during salivary gland development is required in future experiments.

Together, our study has demonstrated the Foxa2 lineage as a critical lineage for salivary gland primordial formation and for generating fully matured, whole salivary glands via CBC, which will serve as a crucial experimental basis for future interspecies blastocyst complementation using human iPSC^43–45^.

## Supplementary Figure Legends

**Supplementary Figure 1. Lineage-tracing, Fgfr2 loss of function, and scRNA-seq analysis of Cre driver candidate genes for CBC**

**(A)** Representative confocal imaging of the salivary gland in the E18.5 *Shh^Cre/+^; Rosa^LSL-tdTomato/+^* (left) and *Shh^Cre/+^; Fgfr2^flox/flox^; Rosa^LSL-tdTomato/+^* mouse (right). Immunostaining of tdTomato: red, E-cadherin (E-cad): cyan, DAPI: gray. (N=3) **(B)** Representative confocal imaging of the salivary gland in the E18.5 *Pitx2^Cre/+^; Rosa^LSL-tdTomato/+^* (left) and *Pitx2^Cre/+^; Fgfr2^flox/flox^; Rosa^LSL-tdTomato/+^* mouse (right). Immunostaining of tdTomato: red, E-cadherin (E-cad): cyan, DAPI: gray. (N=3) **(C)** The morphometric analysis: % of tdTomato-positive cells in E-cad^+^ epithelial cells from the E18.5 Shh^Cre/+^; Rosa^LSLtdTomato/+^ or Pitx2^Cre/+^; Rosa^LSLtdTomato/+^ mice. (N = 3 per biological replicates, 5 fields per group). Error bars represent mean ± SD. **(D)** Annotated uniform manifold approximation projection (UMAP) plots of mouse oral epithelium at E12.0. Clusters assigned: ADL, anterodorsal-lateral; ADM, anterodorsal-medial; AVL, anteroventral-lateral; AVM, anteroventral-medial; Di, diastema; IK, initiation knot; In, incisor; Mo, molar; PL, posterior-lateral; PM, posterior-medial; P/S, periderm and suprabasal cells; SG, salivary gland; T, tongue; V1, ventral 1; V2, ventral 2. **(E)** Feature plots for the expression of candidate Cre-driver genes in mouse oral epithelium at E12.0. **(F)** Dot plots for top 10 differentially expressed genes of tongue (T) and posterior-lateral (PL) culsters. **(G)** Annotated UMAP plots of E8.5 mouse embryos. **(H)** Feature plots for the expression of candidate Cre-driver genes in E8.5 mouse embryos. Red dot aria is the boundary region of the oral cavity. **(I)** Representative confocal imaging of the oral cavity in the E9.5 *Shh^Cre/+^; Rosa^LSL-tdTomato/+^* (left) and E9.5 *Pitx2^Cre/+^; Rosa^LSL-tdTomato/+^* mouse (right). Each right panel is an enlarged image of a white dotted box. Shh lineage labeled only the endoderm (Endo) (arrowhead) but not ectoderm (Ecto) (arrow). Pitx2 lineage labeled neither of them. tdTomato: red, DAPI: gray. (N=3) Scale bars: 100 μm.

**Supplementary Figure 2. The developing salivary glands rescued by the donor cells had normal salivary gland structure and differentiation**

Representative IF-confocal imaging of complemented salivary gland from the E17.5 chimeric mice of *Fgfr2^cKO^*+nEGFP miPSC. The right panel is an enlarged image of a white dotted box. Immunostaining of tdTomato: red, EGFP: green, each cell type maker: cyan, DAPI: gray. (N=3) Scale bars: 100 μm.

## Materials and Methods

### Mouse

Shh^Cre/+^ mice (cat. 05622), Rosa26^tdTomato/tdTomato^ mice (cat. 07914) were obtained from the Jackson Lab. X. Zhang kindly gifted Fgfr2^flox/flox^ mice. We further backcrossed these mice for over three generations with CD-1 mice (cat. 022) from the Charles River. Dr. Nicole C Dubois kindly provided Foxa2^Cre/Cre^ mice (129xB6 mixed background)^46^. For conditional deletion of Fgfr2 (Fgfr2cKO), we crossed Fgfr2^flox/flox^; Rosa26^tdTomato/tdTomato^ females with Foxa2^Cre/Cre^; Fgfr2^flox/+^, Foxa2^Cre/+^; Fgfr2^flox/+^ or Shh^Cre/+^; Fgfr2^flox/+^ males, respectively. PCR performed genotyping of the Shh-Cre and Rosa26-tdTomato alleles according to the protocol provided by the vendor. For the CBC, the genotyping of chimeric animals was confirmed by GFP-negative sorted liver cells and lung cells. For detecting the Fgfr2 floxed allele, we performed PCR using the primer sets: FR2-F1, 5’ ATAGGAGCAACAGGCGG-3’, and FR2-F2, 5’-CAAGAGGCGACCAGTCA-3’^13^. All animal experiments were approved by Columbia University Institutional Animal Care and Use Committee in accordance with US National Institutes of Health guidelines.

### Culture of mouse iPSCs and PSC

We cultured iPSC in a2i/VPA/LIF medium on a feeder, as previously reported^13^. These PSC cells were passaged at a split ratio 1:10 every 2–3 d. For the CBC donor cell preparation, nGFP+iPSCs, PSC^CAG-EGFP^, cultured in a2i/VPA/LIF ^13^, were trypsinized and resuspended in 4 ml cold DMEM + 10% FBS immediately, and filtering the cells with a 40-μm filter. Cells were centrifuged at 350 rcf, 4 °C, for 3 min, and the supernatant was removed. After being washed with flow buffer containing 0.2% BSA, 1% Glutamax, and 1μM Y27632, 1 million cells were resuspended in 100μl of flow buffer. The following antibodies were added: Epcam-BV421 (1:50), SSEA1-PE (1:50), CD31-APC (1:50), Zombie Aqua Fixable Viability Kit (1:100). Epcam^high^SSEA1^high^CD31^high^ cells were sorted by FACS (SONY MA900) and subsequently prepared for the injection.

### Immunofluorescence

Before the immunostaining, antigen retrieval was performed using Unmasking Solution (Vector Laboratories, H-3300) for 10 min at around 100 °C by microwave. 4-μm tissue sections were incubated with primary antibodies in the buffer of M.O.M. kit (Vector Laboratories, MKB-2213-1) overnight at 4 °C, washed in PBS, and incubated with secondary antibodies conjugated with Alexa488, 567, or 647 (ThermoScientific, 1:400) with DAPI for 1.5 h, and mounted with ProLong Gold antifade reagent (Invitrogen, P36930). The images were captured by a Zeiss confocal 710 microscopy. The antibodies were listed in the Supplemental Table. 1.

### Flow cytometry of chimeric mouse lung and liver for genotyping

Lung and liver tissues were minced with micro scissors, and 1 ml of pre-warmed dissociation buffer (1 mg/ml DNase (Sigma, DN25), 5 mg/ml collagen (Roche, 10103578001), and 15 U/ml Dispase II (Stemcell Technologies, 7913) in HBSS), incubated at 37 °C on the rocker with 50 r.p.m. speed, and neutralized with the dissociation buffer by FACS buffer containing 2% FBS, Glutamax, 2mM EDTA and 10mM HEPES in HBSS after the 30 min incubation. After filtrating the cells with a 40-μm filter (FALCON, 352235), cell pellets were resuspended with 1 ml of cold RBC lysis buffer (Biolegend, 420301) to lyse the remaining erythrocytes for 5 min on ice and neutralized by 1 ml cold FACS buffer. After that, it was centrifuged them at 350 rcf, 4 °C, for 3 min to remove the lysed blood cells. Cells were resuspended in 500 μl FACS buffer with PI for the subsequent analyses using SONY MA900.

### Blastocyst preparation and embryo transfer

Blastocysts were prepared by mating Foxa2^Cre/Cre^; Fgfr2^flox/+^, Foxa2^Cre/+^; Fgfr2^flox/+^ or Shh^Cre/+^; Fgfr2^flox/+^ males (all 129 x B6 x CD-1 background) with superovulated Fgfr2^flox/flox^; Rosa26^tdTomato/tdTomato^ females (129 x B6 x CD-1 background). Blastocysts were harvested at E3.5 after superovulation^13^. 20 sorted nGFP^+^ iPSCs were injected into each blastocyst. After the iPSC injection, blastocysts were cultured in an M2 medium (Cosmobio) for a few hours in a 37 °C, 5% CO2 incubator for recovery. Then, blastocysts were transferred to the uterus of the pseudopregnant foster mother.

### Statistical analysis

Data analysis was performed using Prism 8. Data acquired by performing biological replicas of two or three independent experiments are presented as the mean ± SD. Statistical significance was determined using a student t-test. *P < 0.05, ns: non-significant.

## Acknowledgments

We thank Zurab Ninish for his technical assistance. We sincerely appreciate the generous support from Dr. Hiromitsu Nakauchi at Stanford University and the considerate support and scientific input from Dr. Wellington Cardoso at the Columbia Center for Human Development (CCHD) and the members of Cardoso’s lab and CCHD. We acknowledge the support from the CCHD Medicine Microscopy core (MMC), Columbia Stem Cell Initiative (CSCI) Flow Cytometry core (SONY MA900), and Genetically Modified Mouse Model Shared Resource (GMMMSR) for blastocyst injection. We thank Dr. Heiko Lickert of the Technical University of Munich (TUM) for sharing Foxa2-Cre mice. This work was funded by NIH-NHLBI 1R01 HL148223-01, DoD PR190557, PR191133 to M. M., JSPS21KK0290, and The Uehara Memorial Foundation to J. T.

## Notes

### Competing Interest Statement

The authors have declared no competing interest.

