## Supplementary figures and images for "Generation of salivary glands derived from pluripotent stem cells via conditional blastocyst complementation"

### Supplementary Figure

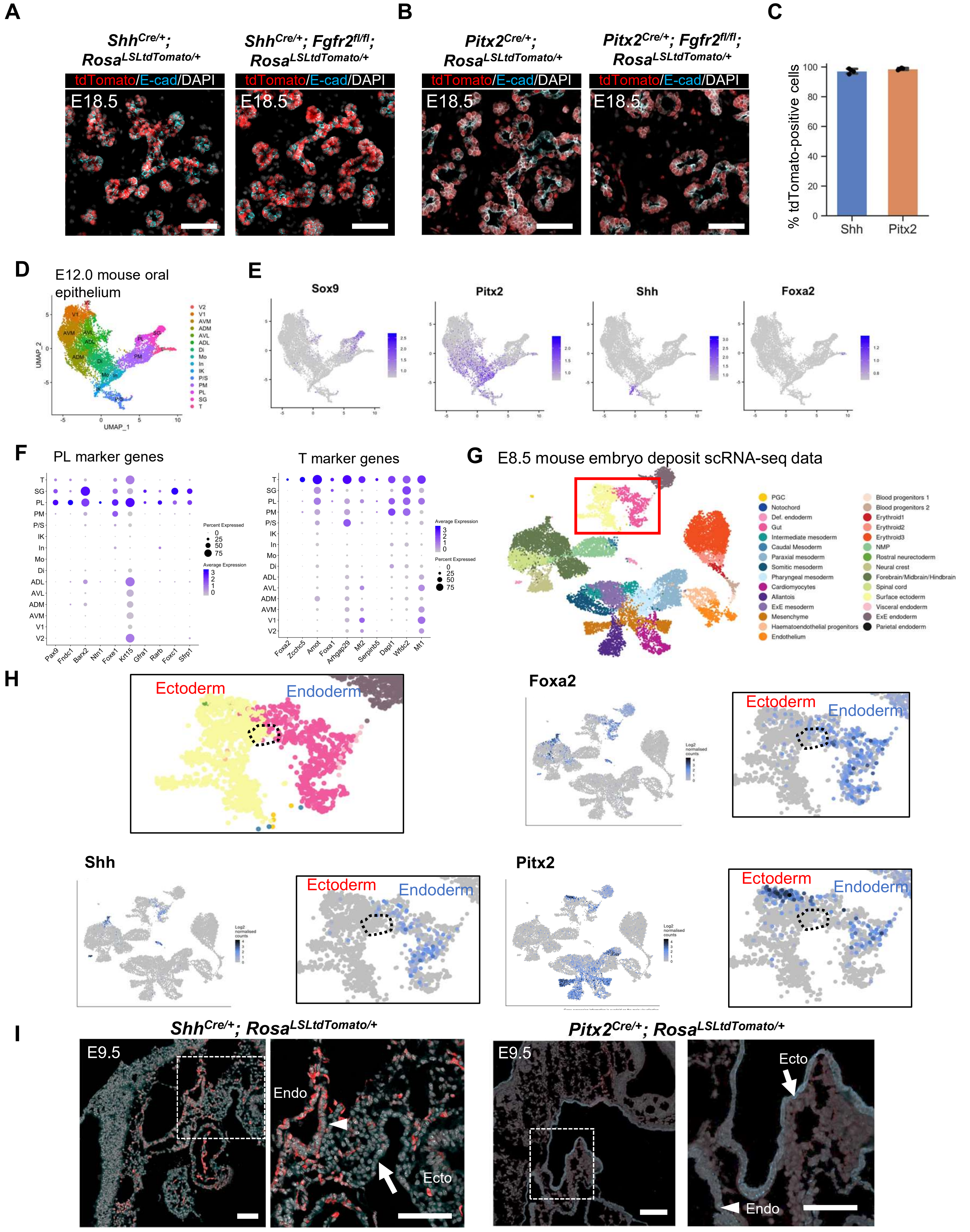

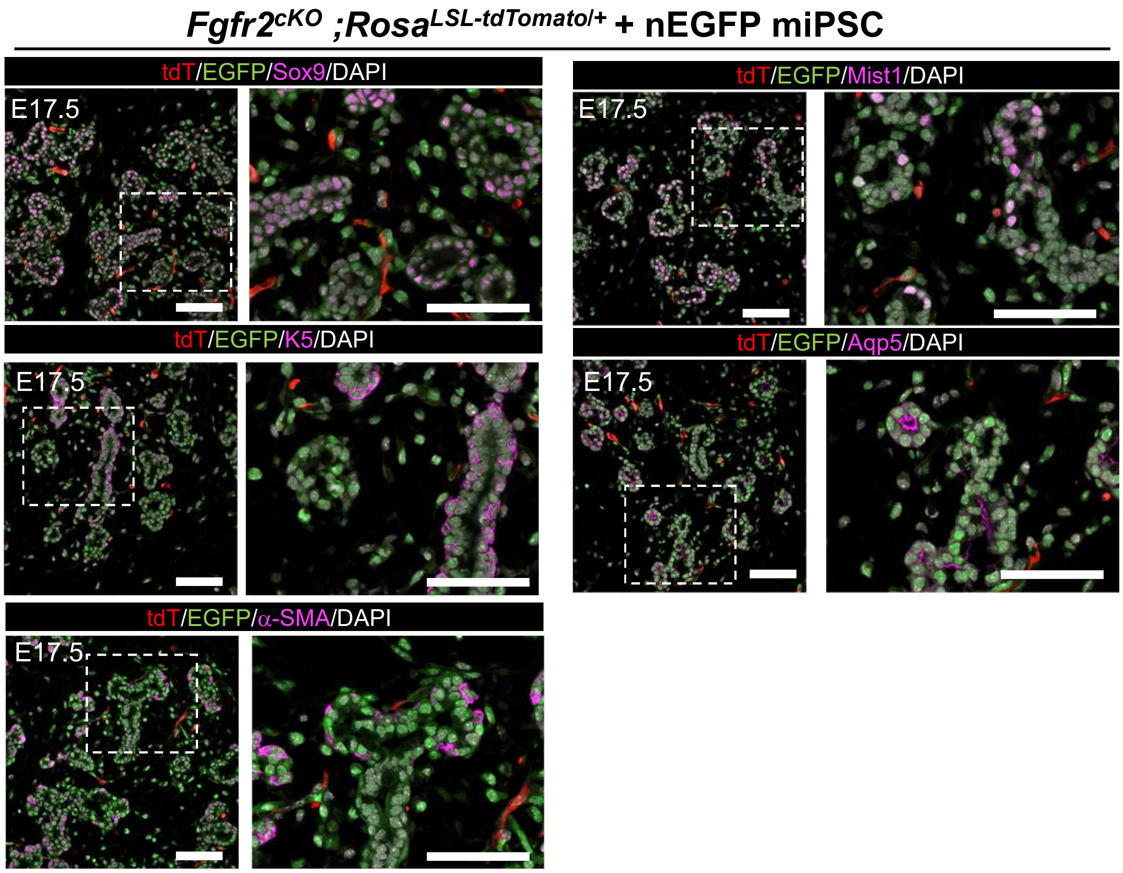
