## Supplementary Table 1 for "Generation of salivary glands derived from pluripotent stem cells via conditional blastocyst complementation"

Table S1

| REAGENT or RESOURCE | SOURCE | IDENTIFIER |
|---|---|---|
| Antibodies | | |
| Anti-E-cadherin | Invitrogen | 131900 |
| Anti-tdTomato | Abcam | Orb182397 |
| Anti-aSMA | Sigma-Aldrich | Ab5694 |
| Anti-Foxa2 | Cell Signaling Technology | 8186S |
| Anti-Sox9 | Sigma-Aldrich | AB5535 |
| Anti-GFP | Aves lab | GFP1020 |
| Anti-Pan-CK | Sigma-Aldrich | C2562 |
| Anti-Mist1 | Cell Signaling Technology | 14896 |
| Anti-CK5 | Abcam | Ab52635 |
| Anti-CK14 | Abcam | Ab7800 |
| Anti-AQP5 | Alomone labs | AQP-005 |
| Donley anti Rabbit IgG (H+L), Alexa Fluor 488 | Invitrogen | A21206 |
| Donley anti Mouse IgG (H+L), Alexa Fluor 488 | Invitrogen | A21202 |
| Donley anti Chicken IgG (H+L), Alexa Fluor 488 | Jackson Immunoresearch | 703545155 |
| Donley anti Rabbit IgG (H+L), Alexa Fluor 568 | Invitrogen | A10042 |
| Donley anti Goat IgG (H+L), Alexa Fluor 568 | Invitrogen | A11057 |
| Donley anti Rabbit IgG (H+L), Alexa Fluor 647 | Invitrogen | A31572 |
| Donley anti Goat IgG (H+L), Alexa Fluor 647 | Invitrogen | A21447 |
